# Mechanistic Modeling of Periodontal Pathogen *Fusobacterium nucleatum*-Induced Epithelial Oncogenic Transformation in Carcinomas via Selected FadA-Lactate-Angiogenesis Pathway

**DOI:** 10.1101/2025.03.10.642351

**Authors:** Timothy Windham, Cris Mantilla, Jimena Garcia

**Affiliations:** University of Houston; University of Houston System at Cinco Ranch: University of Houston

## Abstract

Periodontal pathogens, commonly found in the normal oral flora, have been implicated in cancer progression, yet their metabolic influence remains underexplored. This study investigates the role of *Fusobacterium nucleatum* in tumorigenesis by modeling bacterial-induced inflammation and metabolic shifts. Using computational simulations, we analyze key enzymatic fluxes and their impact on cancer development. Our findings highlight significant lactate accumulation and subsequent VEGF upregulation, suggesting potential biomarkers for early detection and novel therapeutic targets.

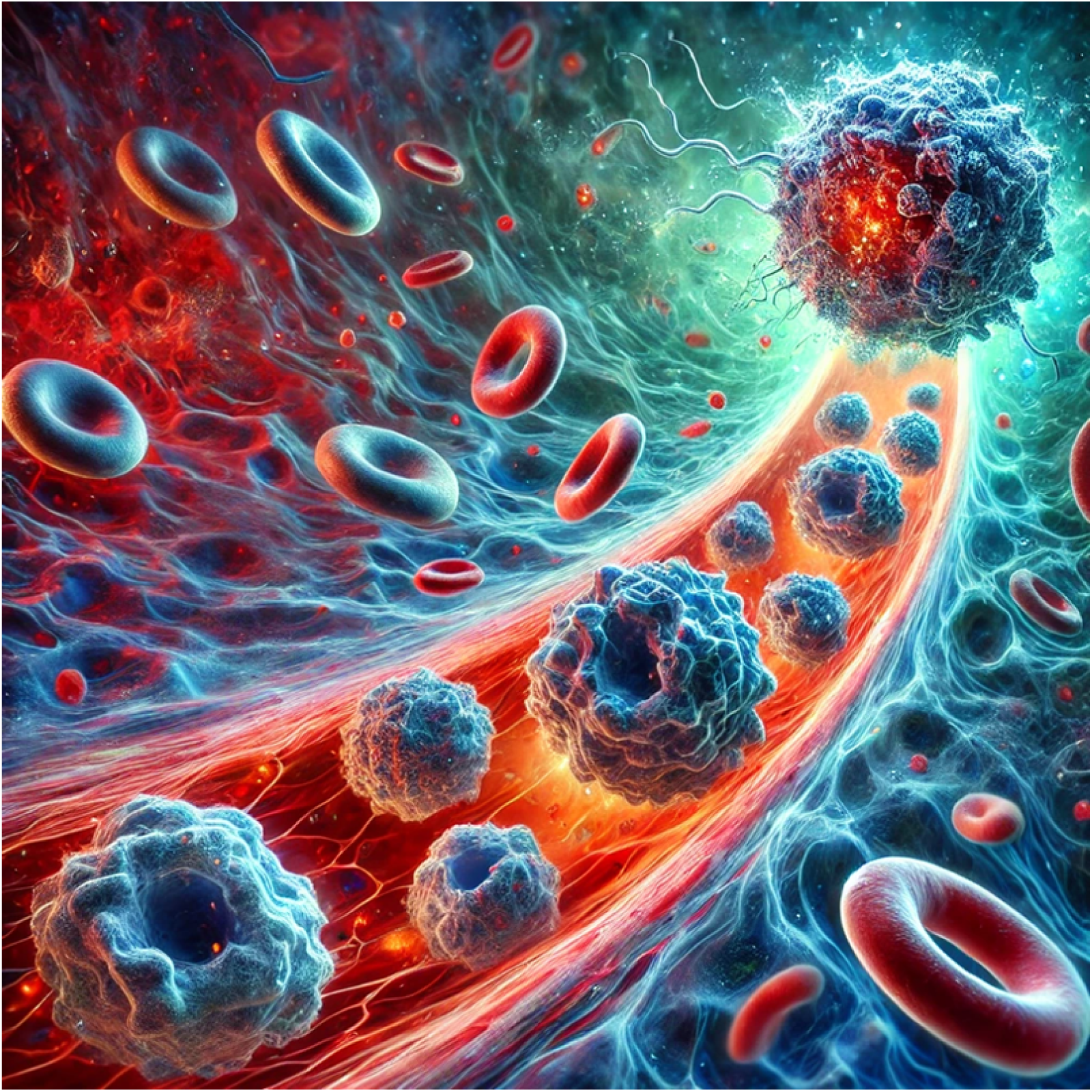

## Introduction and Background

Periodontitis is a chronic inflammatory condition of the gums driven primarily by pathogenic bacteria residing in subgingival biofilms (1). Among these pathogens, *Fusobacterium nucleatum* has garnered significant attention due to its ability to translocate from inflamed gum tissue into systemic circulation and its association with various cancers—most notably colorectal carcinoma (2, 3). Although multiple biochemical pathways have been implicated in linking periodontitis to tumorigenesis, our study focuses on a specific, well-defined pathway to ensure clarity and tractability in our modeling efforts.

In the pathway under investigation, the virulence factor FadA, produced by *F. nucleatum*, binds to the host epithelial cell receptor E-cadherin. This binding event—modeled as a reversible reaction—triggers the activation of intracellular β-catenin, which subsequently transactivates the c-Myc gene, leading to increased synthesis of c-Myc protein. Elevated c-Myc levels drive further oncogenic processes, such as the upregulation of lactate dehydrogenase A (LDHA). The LDHA enzyme catalyzes the conversion of pyruvate to lactate; the resulting lactate accumulation then inhibits prolyl hydroxylase (PHD), stabilizing and activating HIF-1α. Activated HIF-1α induces the expression of vascular endothelial growth factor (VEGF), thereby promoting angiogenesis and facilitating tumor cell proliferation (4, 5).

We specifically selected the FadA–E-cadherin–lactate–angiogenesis pathway for investigation due to its clearly defined molecular interactions and recent identification as a critical mechanistic link between periodontal pathogens and epithelial carcinogenesis. Compared to broader and less well-characterized inflammatory or genetic pathways, this route provides precise, quantifiable targets ideal for both computational modeling and potential therapeutic intervention.

By quantitatively modeling measurable components such as bacterial load, protein concentrations, and cytokine levels, and employing literature-derived and in vivo–approximated kinetic rate constants, we simulate the dynamic behavior of this pathway under both healthy and disease states. This targeted computational approach simplifies our analysis and generates clear insights into potential therapeutic opportunities within this pivotal oncogenic pathway (6, 7).

## Methods for Model Construction

### Reactions Included and Excluded

The following biochemical reactions (detailed in **Figure 1**) were explicitly incorporated into our model:

- Reversible binding of *Fusobacterium nucleatum* virulence factor FadA to epithelial E-cadherin receptors, initiating intracellular signaling cascades (4,5).
- Reversible activation of E-cadherin and subsequent activation and nuclear translocation of β-catenin, enabling transcriptional activity (4,8).
- β-catenin-mediated transcriptional activation of the c-Myc gene, leading to increased production of c-Myc protein, a critical oncogenic factor driving cellular proliferation (9,10).
- c-Myc-driven upregulation of lactate dehydrogenase A (LDHA), facilitating the conversion of pyruvate into lactate, characteristic of the Warburg effect common in cancer metabolism (10,11).
- Lactate-induced inhibition of prolyl hydroxylase (PHD), stabilizing hypoxia-inducible factor-1 alpha (HIF-1α) and leading to its transcriptional activation (6,7).
- HIF-1α-induced upregulation of vascular endothelial growth factor (VEGF), promoting angiogenesis and nutrient supply to support tumor growth (12,13).

**Figure 1.**
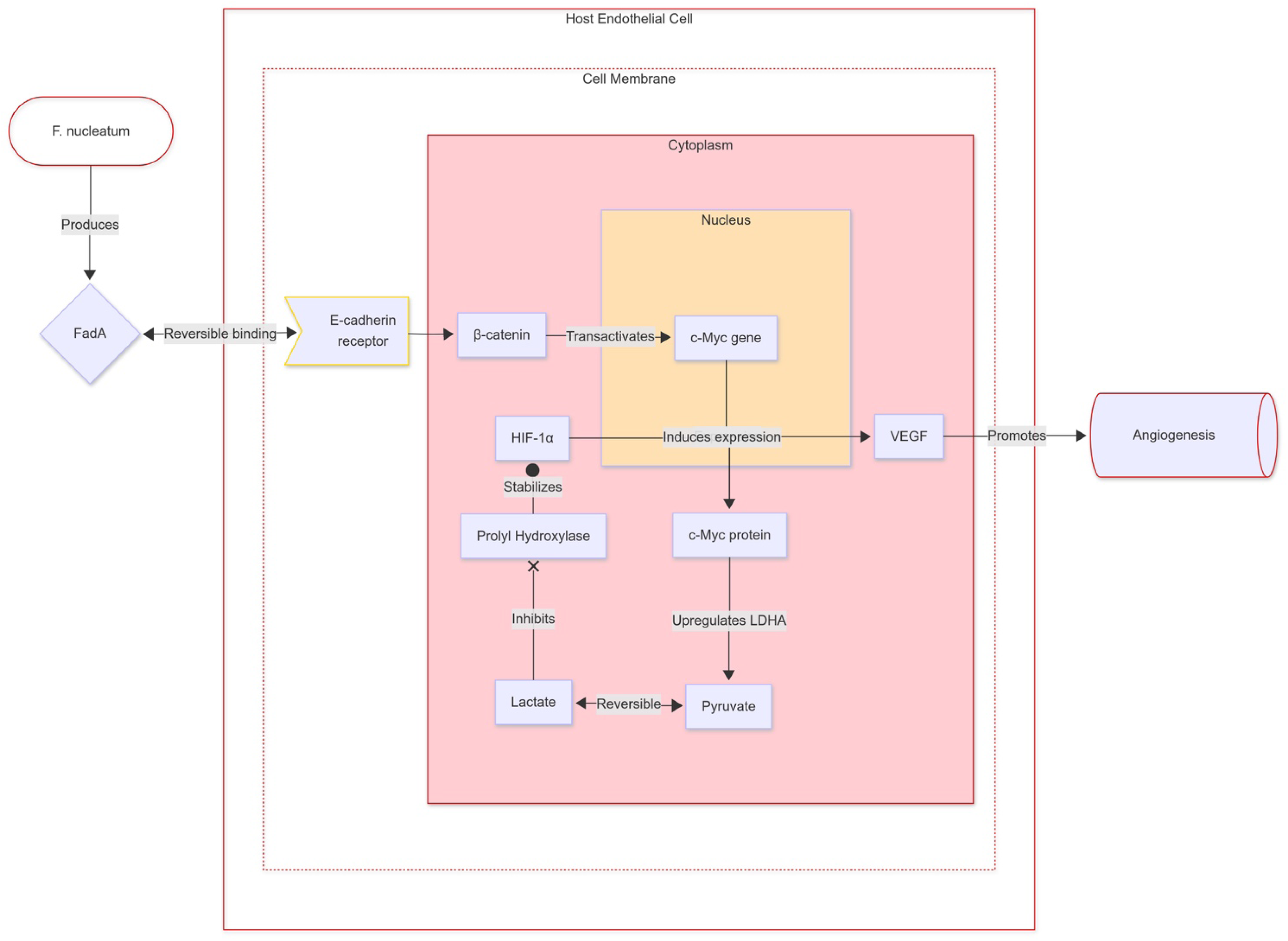
Mechanistic Diagram of the FadA–E-cadherin–Lactate– Angiogenesis Pathway in Host Epithelial Cells Induced by Fusobacterium nucleatum. This figure illustrates the biochemical signaling cascade initiated by the periodontal pathogen *Fusobacterium nucleatum*. The pathogen-derived adhesin FadA binds reversibly to host cell E-cadherin receptors, activating intracellular β-catenin signaling. Activated β-catenin translocates to the nucleus, transactivating the c-Myc gene. Increased c-Myc protein expression subsequently upregulates lactate dehydrogenase A (LDHA), catalyzing the reversible conversion of pyruvate to lactate. Lactate accumulation inhibits prolyl hydroxylase (PHD), stabilizing hypoxia-inducible factor 1-alpha (HIF-1α). Stabilized HIF-1α enhances the expression of vascular endothelial growth factor (VEGF), promoting angiogenesis and potentially facilitating carcinogenesis.

To maintain clarity and computational tractability, we intentionally excluded the following alternative pathways linking periodontitis to tumorigenesis to keep the focus on the selected lactate-angiogenesis axis (3).

- Cytokine-driven immune modulation
- Direct microbial DNA integration
- Regulation of cell cycle progression via Cyclin D1-CDK4/6 and Cyclin E-CDK2 complexes, essential for uncontrolled epithelial proliferation and tumorigenesis (14,15).

### Model Assumptions

Several critical assumptions simplified the model construction:

1. **Constant Pathogen Load:** The concentration of *F. nucleatum* was assumed constant, reflecting chronic infection typically observed in periodontitis (1,3).
2. **Steady-State Conditions:** We assumed that intermediate enzyme-substrate complexes rapidly reached steady-state equilibria, permitting simplified kinetic modeling (17,18).
3. **Homogeneous System Assumption:** All reactions were assumed to occur uniformly within a well-mixed cellular environment, neglecting spatial gradients (16,17).

### Stoichiometric Matrix Modeling and Flux Equations

Our extended model employs a stoichiometric matrix approach utilizing MATLAB to quantitatively represent the biochemical interactions within the FadA–E-cadherin–lactate– angiogenesis pathway (16,17,18) Each relevant biochemical species in the pathway was explicitly defined as follows:

- *S*_1_: FadA
- *S*_2_: E-cadherin
- *S*_3_: FadA–E-cadherin complex
- *S*_4_: β-catenin
- *S*_5_: c-Myc
- *S*_6_: LDHA
- *S*_7_: Pyruvate
- *S*_8_: Lactate
- *S*_9_: HIF-1α
- *S*_10_: VEGF

The reactions that form our pathway are defined by their biochemical transformations, incorporating both reversible and irreversible events as follows:

- *R*_1_: FadA + E-cadherin ⇌ FadA–E-cadherin complex
- *R*_2_: FadA–E-cadherin complex → β-catenin
- *R*_3_: β-catenin → c-Myc
- *R*_4_: c-Myc → LDHA
- *R*_5_: Pyruvate ⇌ Lactate (catalyzed by LDHA)
- *R*_6_: Lactate → HIF-1α (stabilization/activation)
- *R*_7_: HIF-1α → VEGF

The stoichiometric matrix (**Table 1**) constructed from these species and reactions is as follows, where rows correspond to species and columns to reactions, with stoichiometric coefficients indicating the consumption (negative) or production (positive) of species:

**Table 1.** Stoichiometric Matrix representing species production, consumption, and reaction relationships in the FadA–E-cadherin–Lactate–Angiogenesis Pathway. Positive (blue) and negative (red) values indicate species production and consumption, respectively.

| | $R_1$ | $R_2$ | $R_3$ | $R_4$ | $R_5$ | $R_6$ | $R_7$ |
| --- | --- | --- | --- | --- | --- | --- | --- |
| $S_1$ (*FadA*) | (-1) | (0) | (0) | (0) | (0) | (0) | (0) |
| $S_2$ (*E-cadherin*) | (-1) | (0) | (0) | (0) | (0) | (0) | (0) |
| $S_3$ (*FadA-E-cadherin*) | (+1) | (-1) | (0) | (0) | (0) | (0) | (0) |
| $S_4$ (β-catenin) | (0) | (+1) | (-1) | (0) | (0) | (0) | (0) |
| $S_5$ (c-Myc) | (0) | (0) | (+1) | (-1) | (0) | (0) | (0) |
| $S_6$ (LDHA) | (0) | (0) | (0) | (+1) | (0) | (0) | (0) |
| $S_7$ (Pyruvate) | (0) | (0) | (0) | (0) | (-1) | (0) | (0) |
| $S_8$ (Lactate) | (0) | (0) | (0) | (0) | (+1) | (-1) | (0) |
| $S_9$ (HIF-1α) | (0) | (0) | (0) | (0) | (0) | (+1) | (-1) |
| $S_{10}$ (VEGF) | (0) | (0) | (0) | (0) | (0) | (0) | (+1) |

The dynamics of the pathway are described using the following general mass-balance equation:

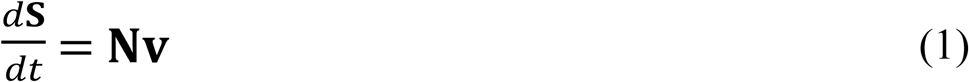

Where **S** is the concentration vector of biochemical species, and **v** is the vector of reaction fluxes defined by kinetic rate expressions. Specifically, we utilized reversible mass-action kinetics for binding interactions, exemplified by the reaction *R*_1_:

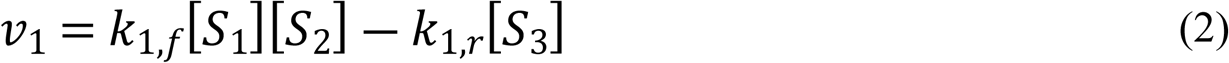

and Michaelis-Menten kinetics for enzymatic conversions, exemplified by the reaction:

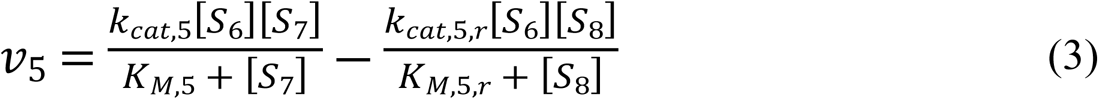

In cases of irreversible enzymatic reactions or simplified transformations, a first-order approximation was utilized:

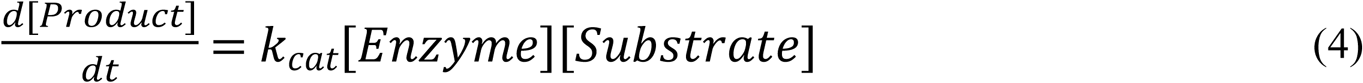

Kinetic rate constants (k-values) were primarily sourced from validated biochemical databases, including BRENDA and KEGG (19,20). When specific experimental kinetic data were unavailable, parameters were approximated from analogous well-characterized enzyme-substrate systems. For example:

- The reversible binding kinetics between FadA and E-cadherin (*k*_1,_*_f_*,*k*_1,*r*_) were approximated from established bacterial adhesin-host receptor interactions (5).
- Kinetic parameters for LDHA-catalyzed pyruvate-lactate interconversion (*K_M_*_5_,*k*_*cat*,5,*r*_,*K*_*M*,5,*r*_) were estimated from established human LDHA activity relevant to the Warburg effect observed in cancer metabolism (10,11).
- Lactate-induced PHD inhibition kinetics (*k*_6_) were approximated from published data on similar metabolic inhibitors (6,7).

These parameter approximations (**Table 2**) were carefully justified through detailed biological rationale and supported by extensive literature review, ensuring the physiological relevance and robustness of our computational simulations (16,17,18,19,20).

**Table 2.** Comprehensive Summary of Enzymes, Reactions, Kinetic Parameters (k-values), and Associated Ordinary Differential Equations (ODEs) in the FadA–E-cadherin–Lactate– Angiogenesis Pathway.

| Reaction | Enzyme/Process | Reaction Description | Kinetic Parameters (k-values) | Associated ODEs and Estimated k-values |
| --- | --- | --- | --- | --- |
| $R_1$ | Adhesin-Receptor Interaction (Reversible) | FadA + E-cadherin ⇌ FadA–E-cadherin complex | $k_{1,f}, k_{1,r}$; approximated from InlA–E-cadherin binding kinetics (5) | $d[S_3]/dt = k_{1,f}[S_1][S_2] - k_{1,r}[S_3]$ $k_{1,f} = 10^6 M^{-1} s^{-1}$ $k_{1,r} = 10^{-2} s^{-1}$ |
| $R_2$ | Signal Transduction | FadA–E-cadherin complex → β-catenin | $k_2$; estimated from β-catenin signaling kinetics (8) | $d[S_4]/dt = k_2[S_3] - k_3[S_4]$ $k_2 = 10^{-3} s^{-1}$ $k_3 = 10^{-4} s^{-1}$ |
| $R_3$ | Gene Activation | β-catenin → c-Myc | $k_3$; approximated from gene expression kinetics (9,10) | $d[S_5]/dt = k_3[S_4] - k_4[S_5]$ $k_4 = 10^{-4} s^{-1}$ |
| $R_4$ | Gene Product Regulation | c-Myc → LDHA | $k_4$; from c-Myc transcriptional activation of LDHA (10,11) | $d[S_6]/dt = k_4[S_5]$ $k_4 = 10^{-4} s^{-1}$ |
| $R_5$ | LDHA (Reversible) | Pyruvate ⇌ Lactate | $k_{cat,5}, K_{M,5}, k_{cat,5,r}, K_{M,5,r}$; human LDHA kinetic parameters (10,11) | $d[S_8]/dt = \frac{k_{cat,5}[S_6][S_7]}{K_{M,5} + [S_7]} - \frac{k_{cat,5,r}[S_6][S_8]}{K_{M,5,r} + [S_8]}$ $k_{cat,5} = 375 s^{-1}, K_{M,5} = 0.3 mM$ $k_{cat,5,r} = 300 s^{-1}, K_{M,5,r} = 4.5 mM$ |
| $R_6$ | PHD Inhibition by Lactate | Lactate → HIF-1α | $k_6$; approximated from lactate-PHD inhibition kinetics (6,7) | $d[S_9]/dt = k_6[S_8] - k_7[S_9]$ $k_6 = 10^{-4} s^{-1}$ |
| $R_7$ | HIF-1α-mediated VEGF Expression | HIF-1α → VEGF | $k_7$; from HIF-1α-dependent VEGF activation (12,13) | $d[S_{10}]/dt = k_7[S_9]$ $k_7 = 10^{-4} s^{-1}$ |

### Results Derived from the Model

To investigate the dynamics of the FadA–E-cadherin–lactate–angiogenesis signaling pathway, computational simulations were conducted using a stoichiometric matrix approach coupled with kinetic flux equations (**Table 2**). The dynamic concentration profiles of all pathway intermediates under healthy conditions are presented in **Figure 2**.

**Figure 2.**
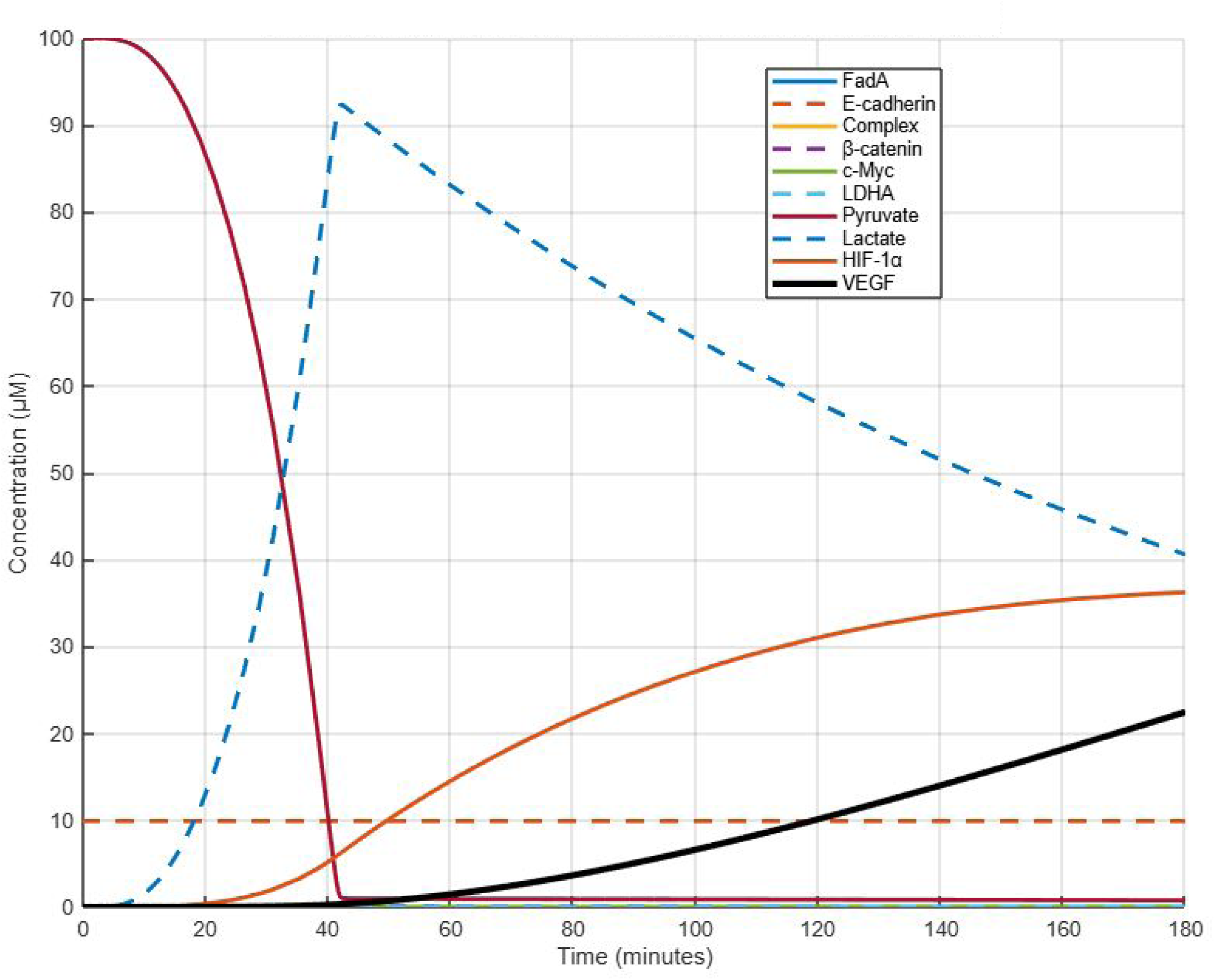
The dynamic concentration profiles of all pathway intermediates under healthy conditions over the 180-minute simulation period. In this scenario, the model predicts a brief transient phase during which substrate concentrations rapidly adjust from their initial values, followed by a gradual progression toward steady state. The resulting conversion of pyruvate to lactate is observed as a steady rise in lactate concentration, which in turn contributes to the stabilization of HIF-1α and the eventual upregulation of VEGF. These changes collectively mirror the expected behavior in a healthy epithelial cellular environment where enzyme kinetics and substrate fluxes balance to maintain homeostasis.

Under periodontitis conditions, simulations started with an elevated initial concentration of FadA (25 nM), representing chronic periodontal infection. As shown in **Figure 3**, this increased pathogen load accelerated the formation of the FadA–E-cadherin complex, amplifying downstream signaling and resulting in enhanced lactate production and robust VEGF expression.

**Figure 3.**
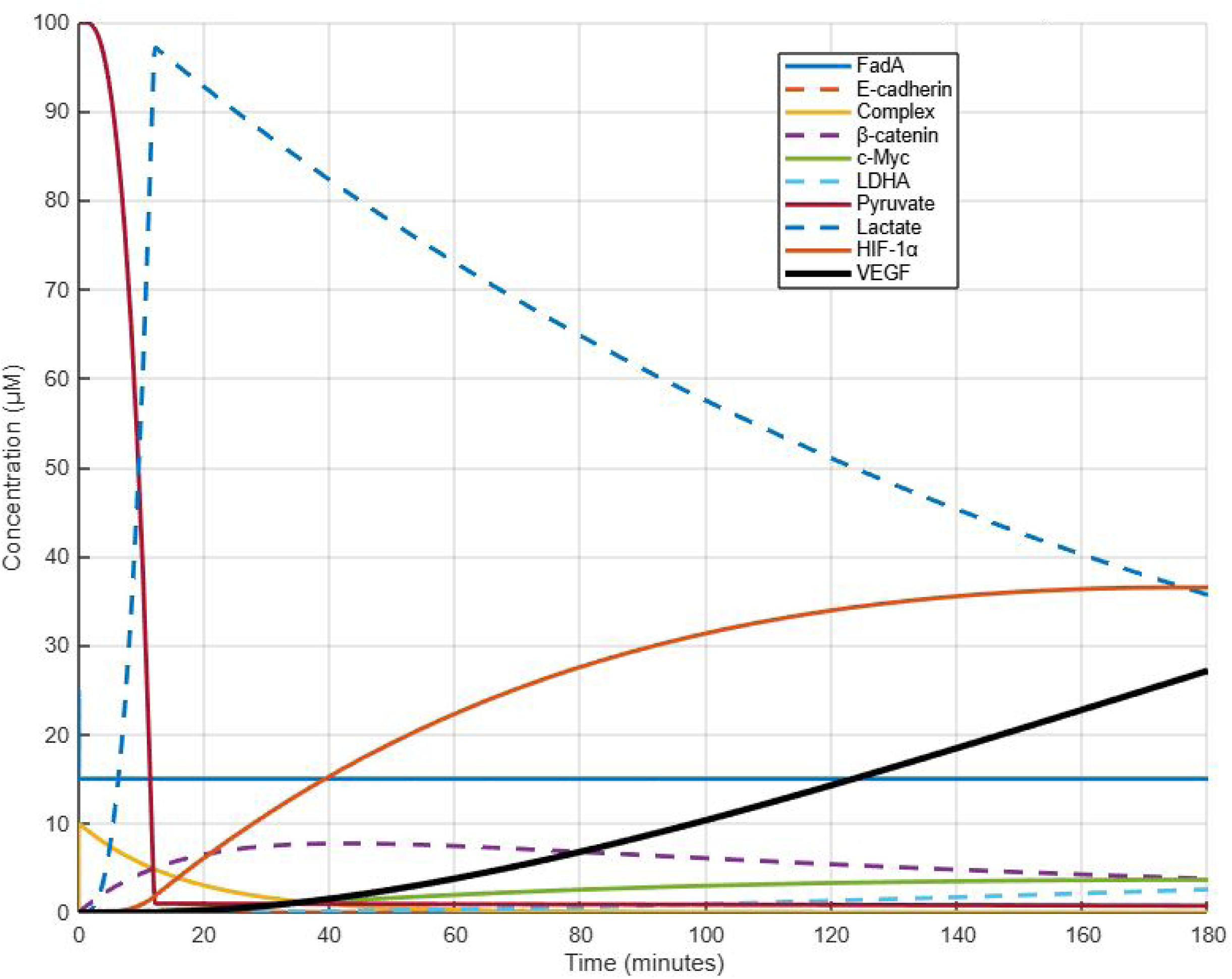
The simulation results for the periodontitis condition, with an elevated initial concentration of FadA (25 nM), representing chronic infection. Under these conditions, the increased FadA accelerates the formation of the FadA–E-cadherin complex and amplifies downstream signaling, rapidly activating β-catenin and inducing c-Myc expression. This heightened metabolic flux results in an earlier and steeper rise in lactate levels, enhanced stabilization of HIF-1α, and robust VEGF expression, indicating significantly intensified pro-angiogenic signals under periodontal disease conditions.

A direct comparison of selected key substrates (FadA, lactate, and VEGF) under healthy versus periodontitis conditions is illustrated in **Figure 4**. These comparisons clearly highlight differential kinetics, confirming the critical influence of *F. nucleatum* on epithelial metabolic reprogramming and angiogenesis.

**Figure 4.**
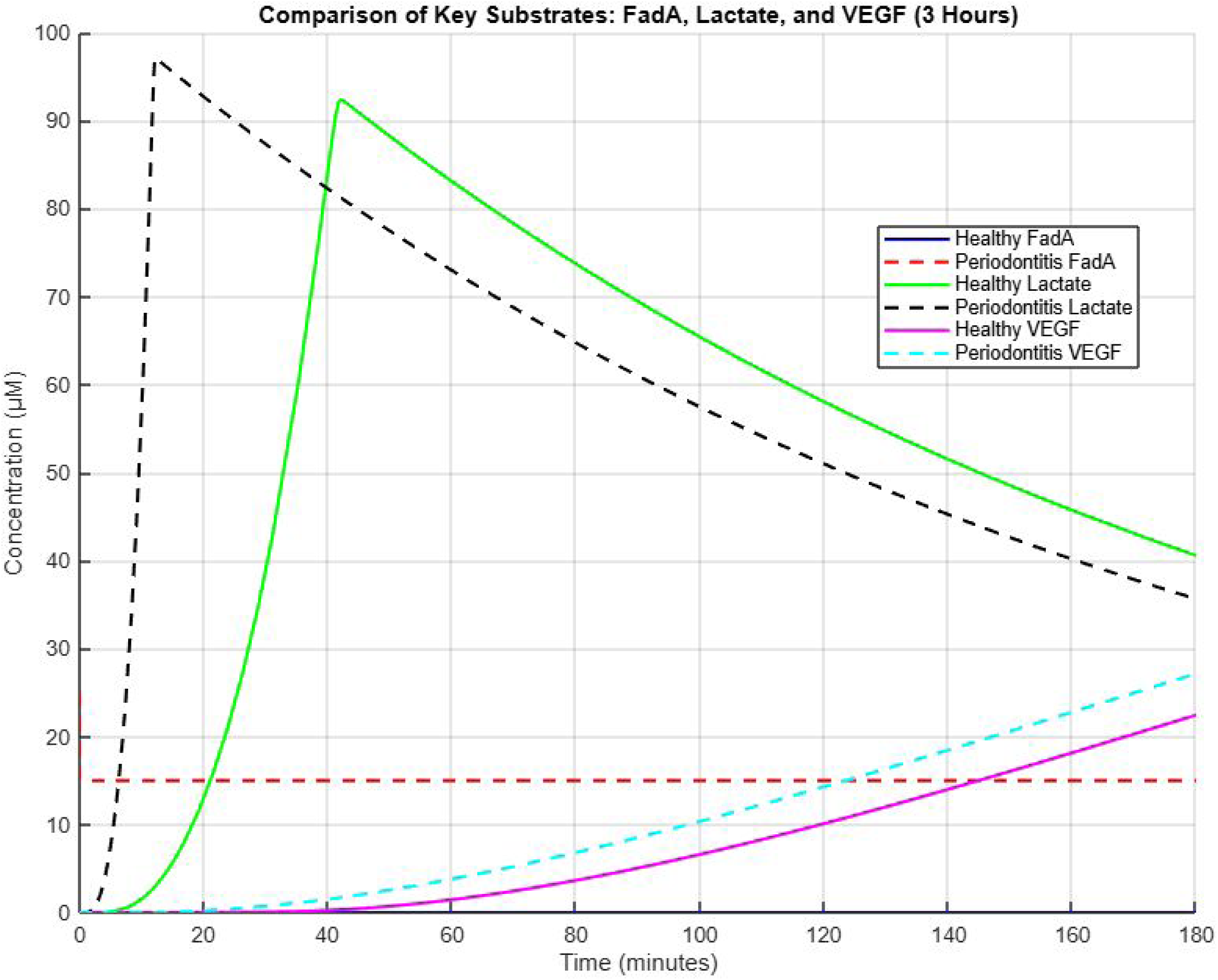
Selected key substrates—specifically FadA, lactate, and VEGF—under healthy and periodontitis conditions, emphasizing differential kinetics. Periodontitis conditions exhibit higher peak concentrations and extended transient phases, confirming the critical influence of *F. nucleatum*-induced alterations in enhancing metabolic reprogramming and angiogenic responses.

To further elucidate individual substrate dynamics, detailed kinetic plots for each biochemical species are provided in Supplementary **Figures S1–S10**.

Overall, the simulation results derived from our stoichiometric matrix approach coupled with kinetic flux equations not only corroborate our mechanistic understanding of the pathway but also offer predictive insights into how alterations in bacterial virulence factors might influence cellular metabolism and angiogenesis. These results will serve as a foundation for further experimental validation and highlight potential targets for therapeutic intervention in cancers associated with periodontal disease.

## Discussion

The computational model simulations highlight the significant role of the FadA–E-cadherin– lactate–angiogenesis signaling pathway in periodontal pathogen-induced cancer progression. **Figure 4** illustrates the predicted amplification of key enzymatic activities—particularly LDHA, HIF-1α, and VEGF—under elevated bacterial loads of *Fusobacterium nucleatum* indicates a clear mechanistic pathway linking periodontitis to increased oncogenic potential. These findings align with previously documented metabolic and inflammatory alterations characteristic of the Warburg effect commonly observed in cancer metabolism (10, 11).

Notably, the enhanced transient dynamics observed under periodontitis conditions support the hypothesis that periodontal pathogens, such as *F. nucleatum*, could significantly intensify pro-angiogenic signals. These results suggest a potential biomarker role for lactate and VEGF levels in early cancer detection and emphasize the critical window for therapeutic intervention targeting early events in the signaling cascade.

Moreover, the incorporation of additional immune and inflammatory pathways in future expanded models could provide deeper insights into the systemic consequences of periodontal infections, enriching our understanding of periodontal pathogen-driven cancer initiation and progression (3). The suggested epidemiological correlation studies would further validate these findings in clinical settings, strengthening the translational relevance of the model predictions.

One limitation of our current model is the assumption of steady-state kinetics and a homogeneous cellular environment, which might not fully capture spatial and temporal complexities observed in biological systems. Addressing these constraints in future studies could enhance the robustness and predictive accuracy of the model.

### Potential Therapeutic and Preventative Targets

Given these mechanistic insights linking *F. nucleatum* to epithelial carcinogenesis, several promising therapeutic and preventive strategies emerge. The proposed therapeutic strategies— including FadA adhesin blockers, LDHA inhibitors, HIF-1α modulators, VEGF antagonists, and targeted probiotic or antimicrobial interventions—highlight tangible avenues for reducing or preventing periodontal pathogen-driven oncogenesis. Preclinical validation and further epidemiological studies will be essential next steps to translate these computational findings into clinical applications effectively.

### FadA Adhesin Blockers

Develop therapeutic agents such as antibodies or small-molecule inhibitors specifically targeting FadA adhesin to block its interaction with E-cadherin. Preventing this initial binding event could effectively disrupt the downstream oncogenic signaling cascade (4, 5).

### LDHA Inhibition

Explore drugs that selectively inhibit LDHA activity, thus decreasing lactate accumulation and subsequent HIF-1α stabilization, potentially reducing angiogenesis and tumorigenesis associated with periodontal pathogen-induced cancer (10, 11).

### HIF-1α Modulators

Evaluate therapeutic strategies targeting HIF-1α stabilization, aiming to control or inhibit VEGF-driven angiogenesis. Such approaches could help mitigate tumor growth and metastatic potential driven by periodontal disease-related inflammation (7, 12).

### VEGF Pathway Antagonists

Utilize existing FDA-approved anti-VEGF drugs (such as bevacizumab) in preclinical studies, assessing their efficacy in preventing or slowing tumor progression in periodontal disease contexts (13).

### Probiotic or Antimicrobial Approaches

Investigate the preventive potential of probiotics or targeted antimicrobials that specifically reduce the prevalence or virulence of *F. nucleatum* in periodontal pockets, thus potentially reducing systemic dissemination and downstream oncogenic effects (3).

### Future Research Directions

Although these therapeutic strategies hold significant potential, further research—including experimental validation and expanded computational modeling—is essential to fully translate these findings into clinical practice.

### Experimental Validation of Key Nodes

To further substantiate computational predictions, experimental validation using in vitro cell culture and animal models is critical. Measuring changes in protein expression levels and enzymatic activities such as LDHA, HIF-1α, and VEGF under varying bacterial loads of *F. nucleatum* will confirm pathway predictions (4, 6, 7, 12).

### Time-Resolved Single-Cell Analyses

Adopting single-cell RNA sequencing or proteomic analyses could provide insights into cell-specific responses to periodontal pathogens. Identifying cellular heterogeneity and potential early biomarkers could significantly enhance early detection strategies for pathogen-driven oncogenesis (8, 2).

### Expanded Pathway Modeling

Extend computational modeling to integrate additional immune-related or inflammatory cytokine-mediated signaling pathways. Such comprehensive modeling would provide a broader view of the systemic impact of periodontal pathogens on cancer initiation and progression (3).

### Clinical Correlation Studies

Perform epidemiological studies correlating periodontal disease severity and pathogen load with cancer incidence or progression rates. Such studies will strengthen the clinical relevance of the model predictions (1, 2).

## Conclusion

In conclusion, this study provides computational evidence supporting the significant impact of the FadA–E-cadherin–lactate–angiogenesis pathway in periodontal pathogen-induced cancer progression. These findings offer clear biomarkers for early detection and identify promising therapeutic targets. Continued experimental validation and expansion of the pathway modeling will further refine our understanding. Importantly, translating these computational insights into clinical practice could lead to novel diagnostic tools and targeted treatments, significantly improving patient outcomes in periodontal disease-associated carcinomas.

---

**Figure S1.**
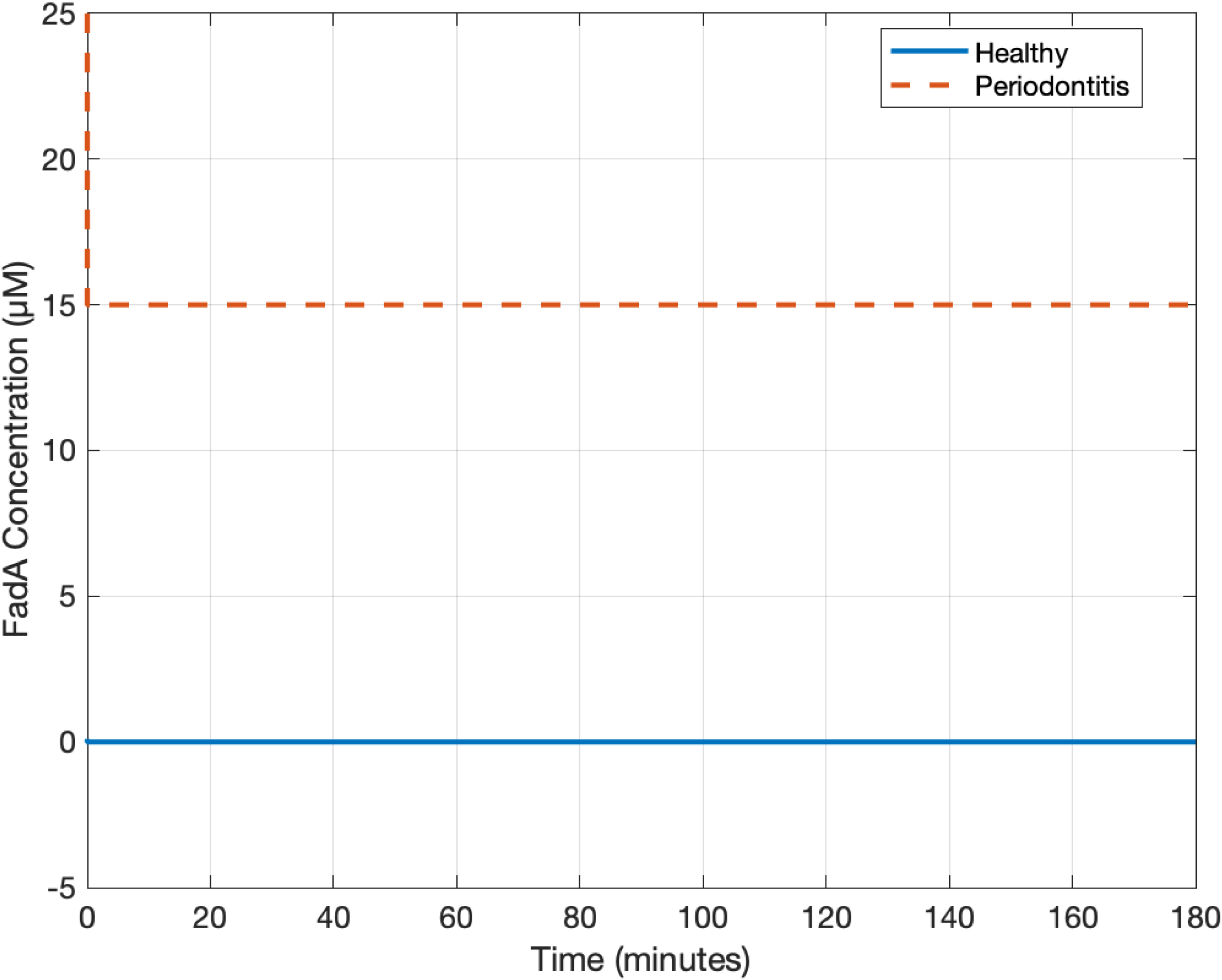
FadA Concentration Dynamics Under Healthy and Periodontitis Conditions. The concentration profile of the FadA virulence factor over three hours, comparing baseline (healthy) conditions to elevated levels during periodontitis.

**Figure S2.**
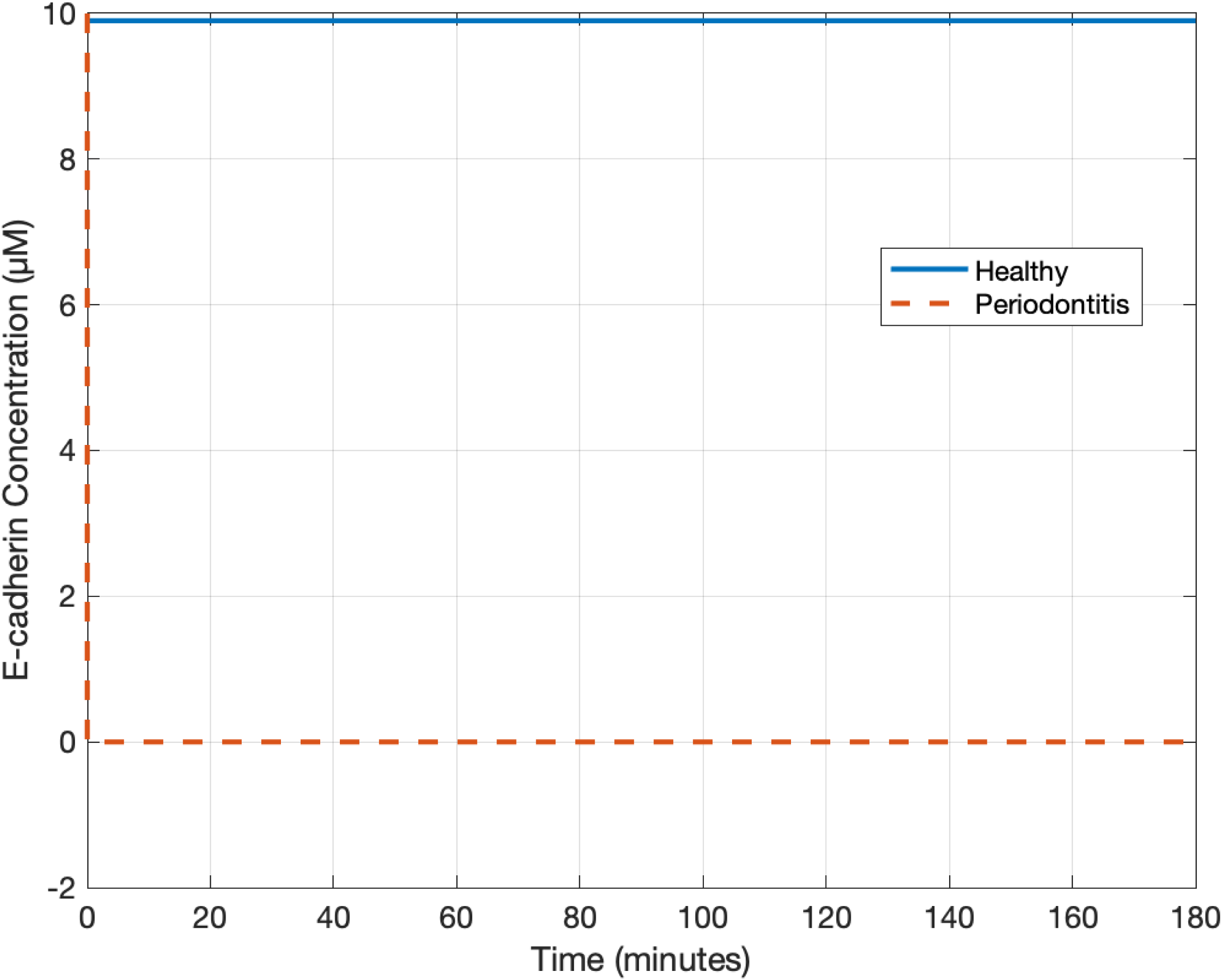
E-cadherin Concentration Dynamics Under Healthy and Periodontitis Conditions. Changes in E-cadherin concentration, highlighting receptor consumption upon increased interaction with FadA under periodontitis conditions.

**Figure S3.**
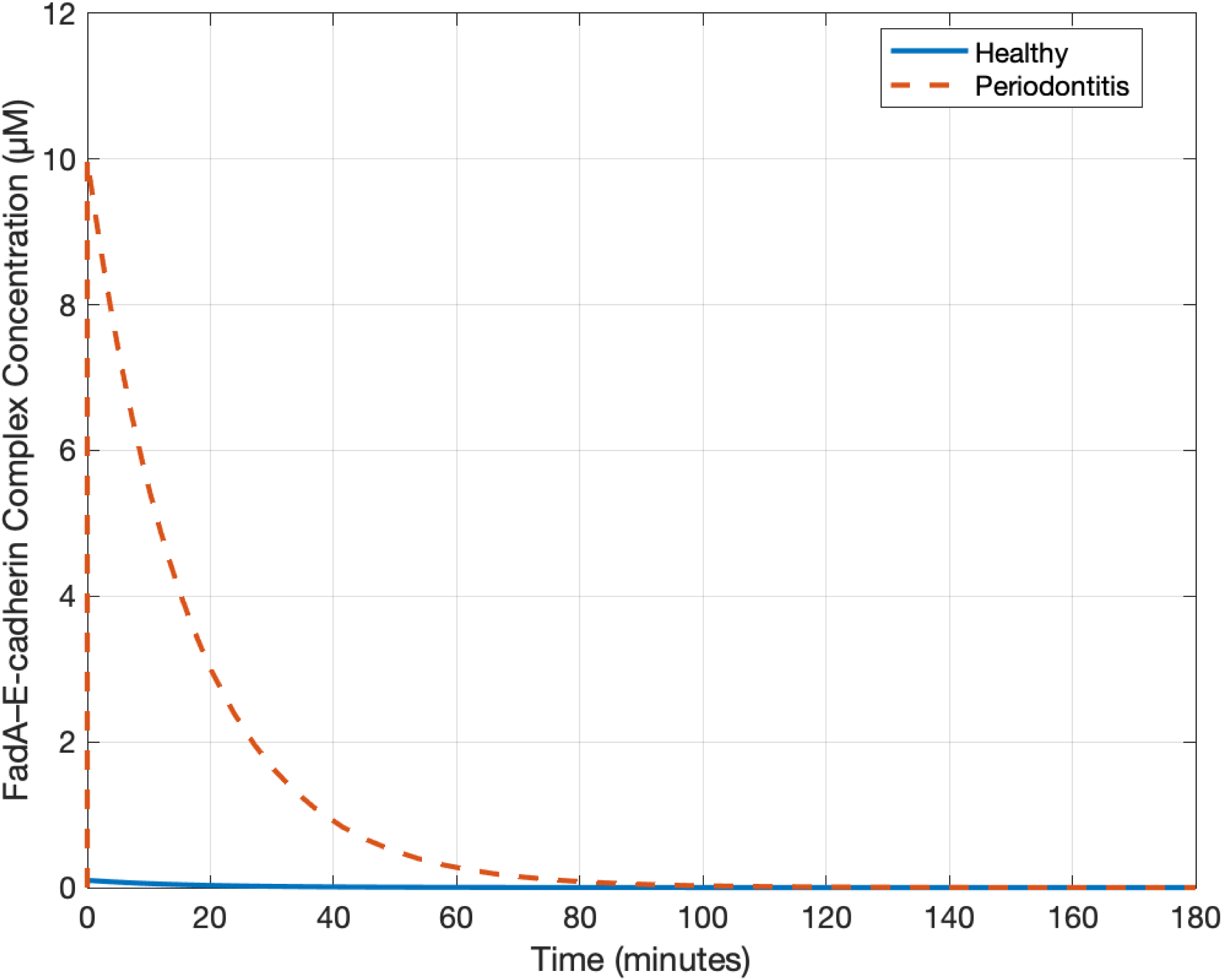
FadA–E-cadherin Complex Dynamics Under Healthy and Periodontitis Conditions. Kinetic profile showing formation of the FadA–E-cadherin complex, with higher complex formation under periodontitis conditions reflecting increased pathogen-host interactions.

**Figure S4.**
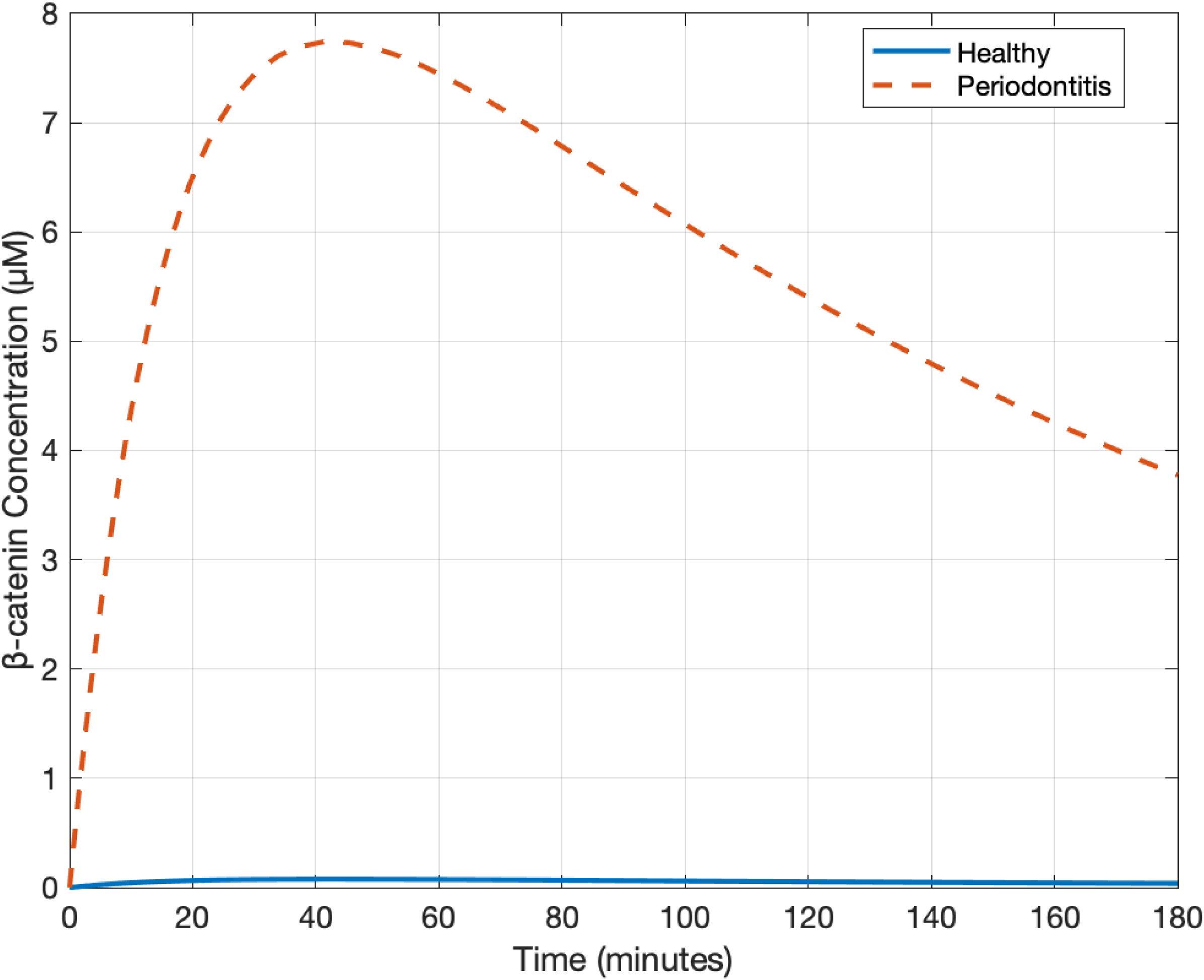
β-catenin Concentration Dynamics Under Healthy and Periodontitis Conditions. Concentration changes of β-catenin, indicating enhanced signaling under chronic periodontal pathogen exposure.

**Figure S5.**
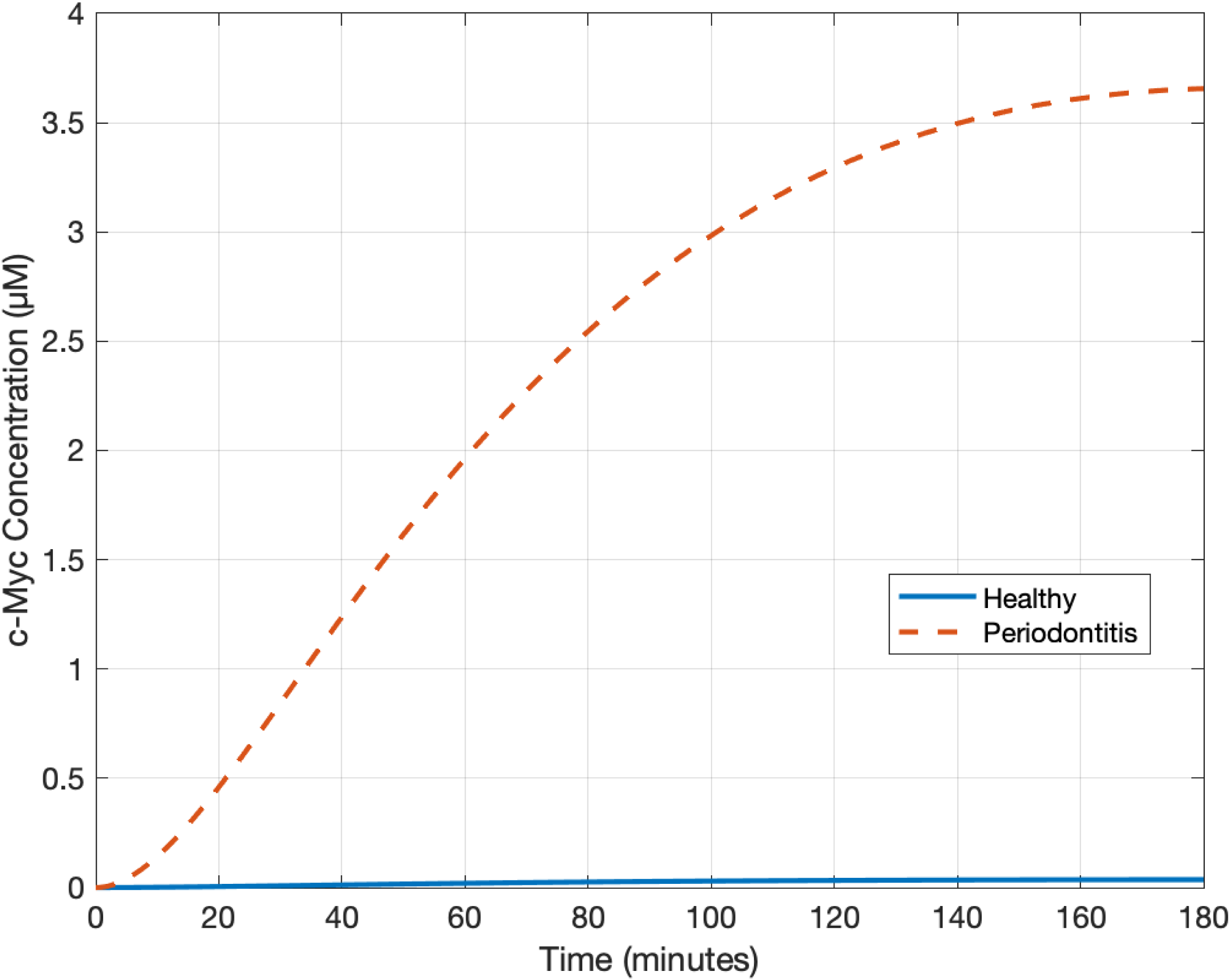
c-Myc Concentration Dynamics Under Healthy and Periodontitis Conditions. Kinetics of c-Myc expression, showing clear oncogenic activation due to periodontal pathogen presence.

**Figure S6.**
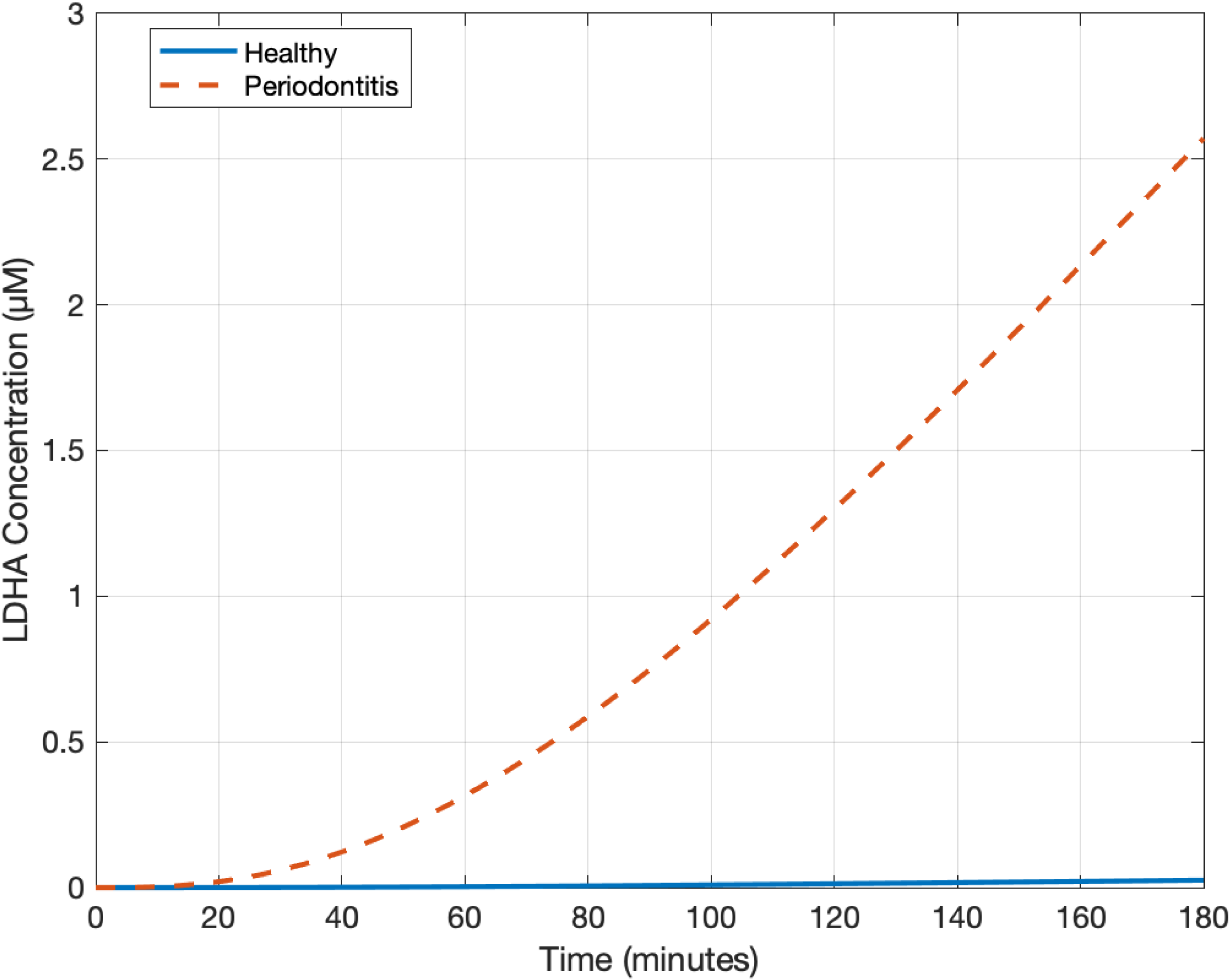
LDHA Concentration Dynamics Under Healthy and Periodontitis Conditions. Levels of lactate dehydrogenase A, critical to lactate production and metabolic shifts commonly associated with cancer cells.

**Figure S7.**
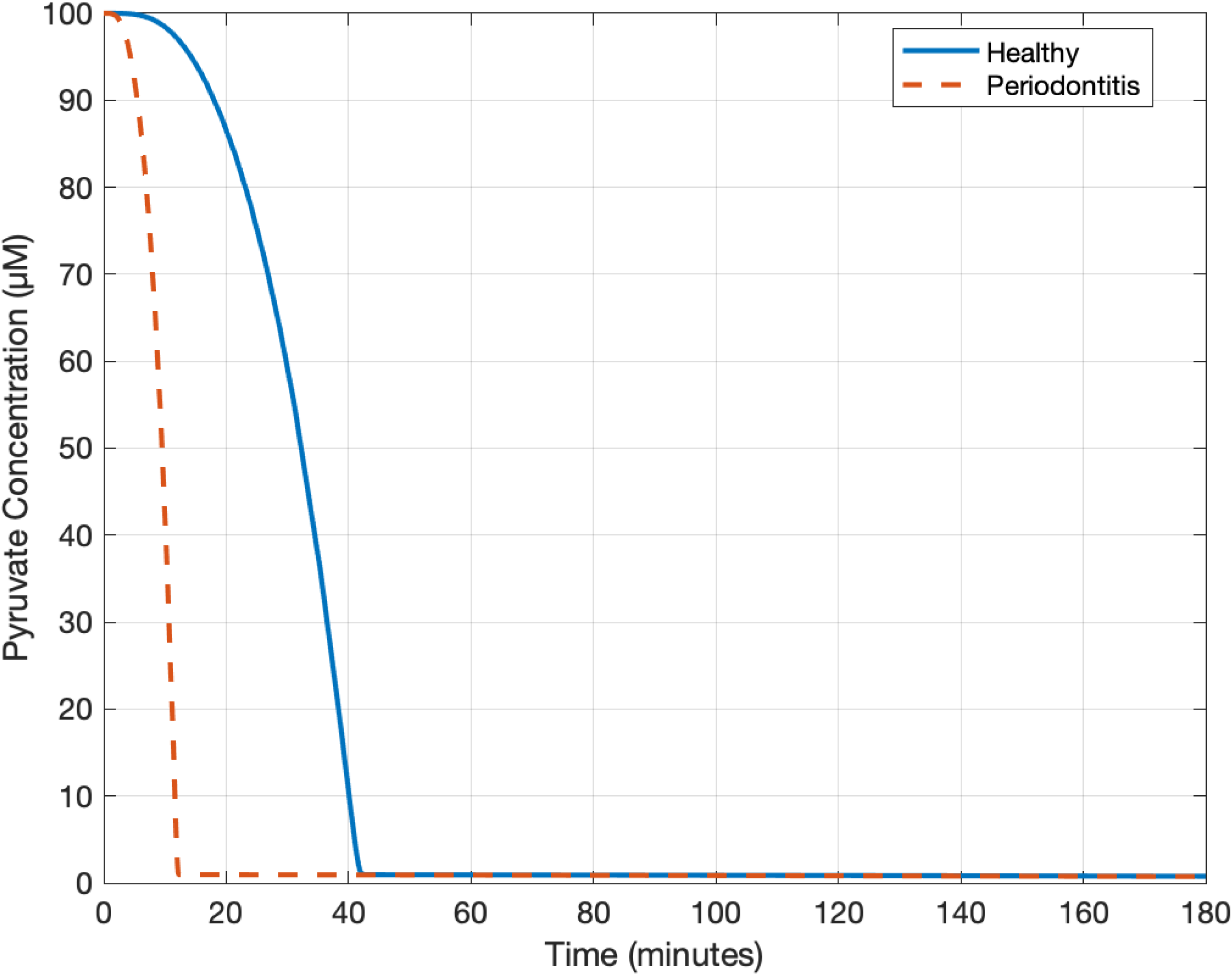
Pyruvate Concentration Dynamics Under Healthy and Periodontitis Conditions. Dynamics of pyruvate, showing the substrate’s consumption pattern altered by increased LDHA activity in disease conditions.

**Figure S8.**
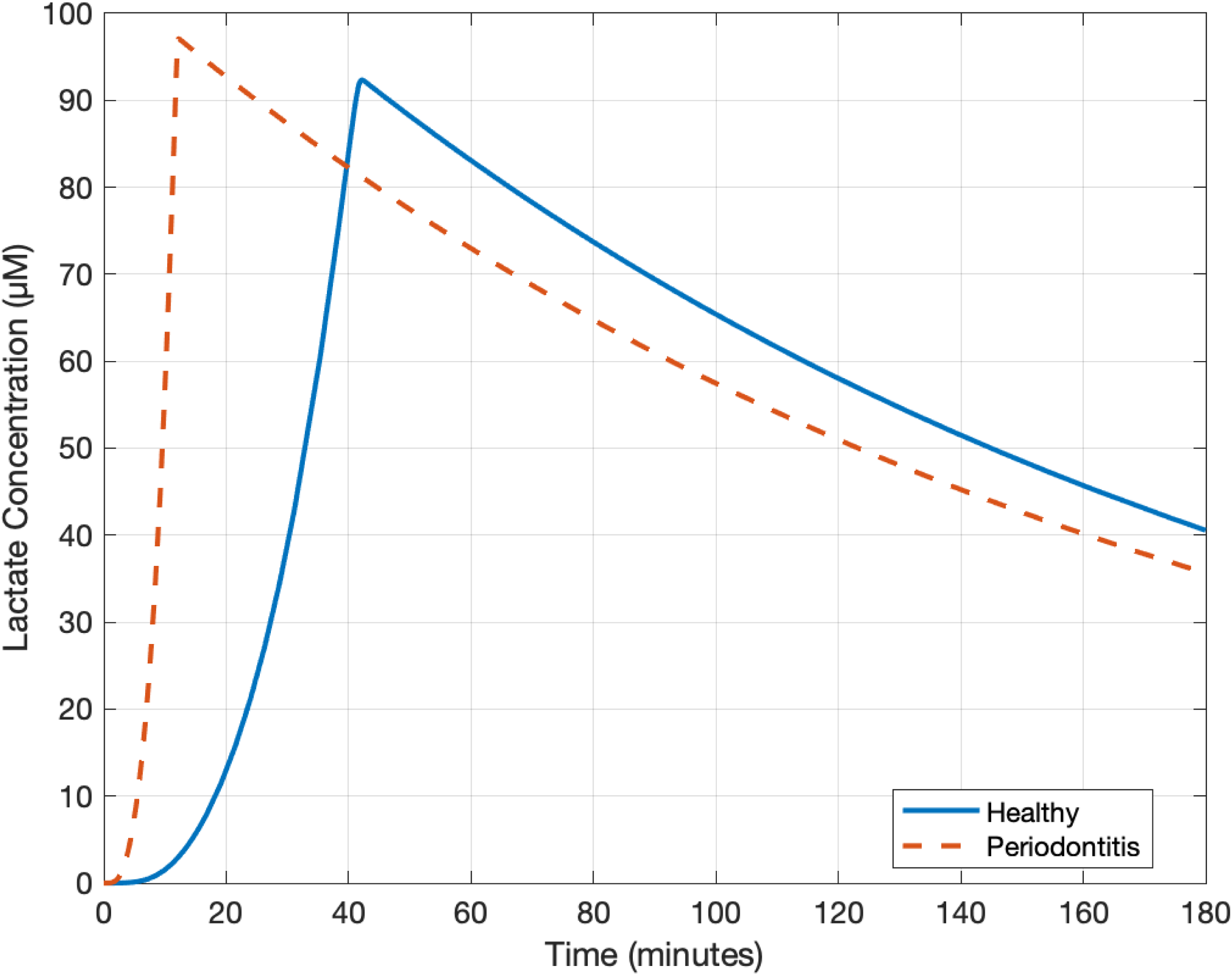
Lactate Concentration Dynamics Under Healthy and Periodontitis Conditions. Simulation highlighting the pronounced lactate accumulation driven by periodontal pathogen-induced metabolic shifts characteristic of tumorigenesis.

**Figure S9.**
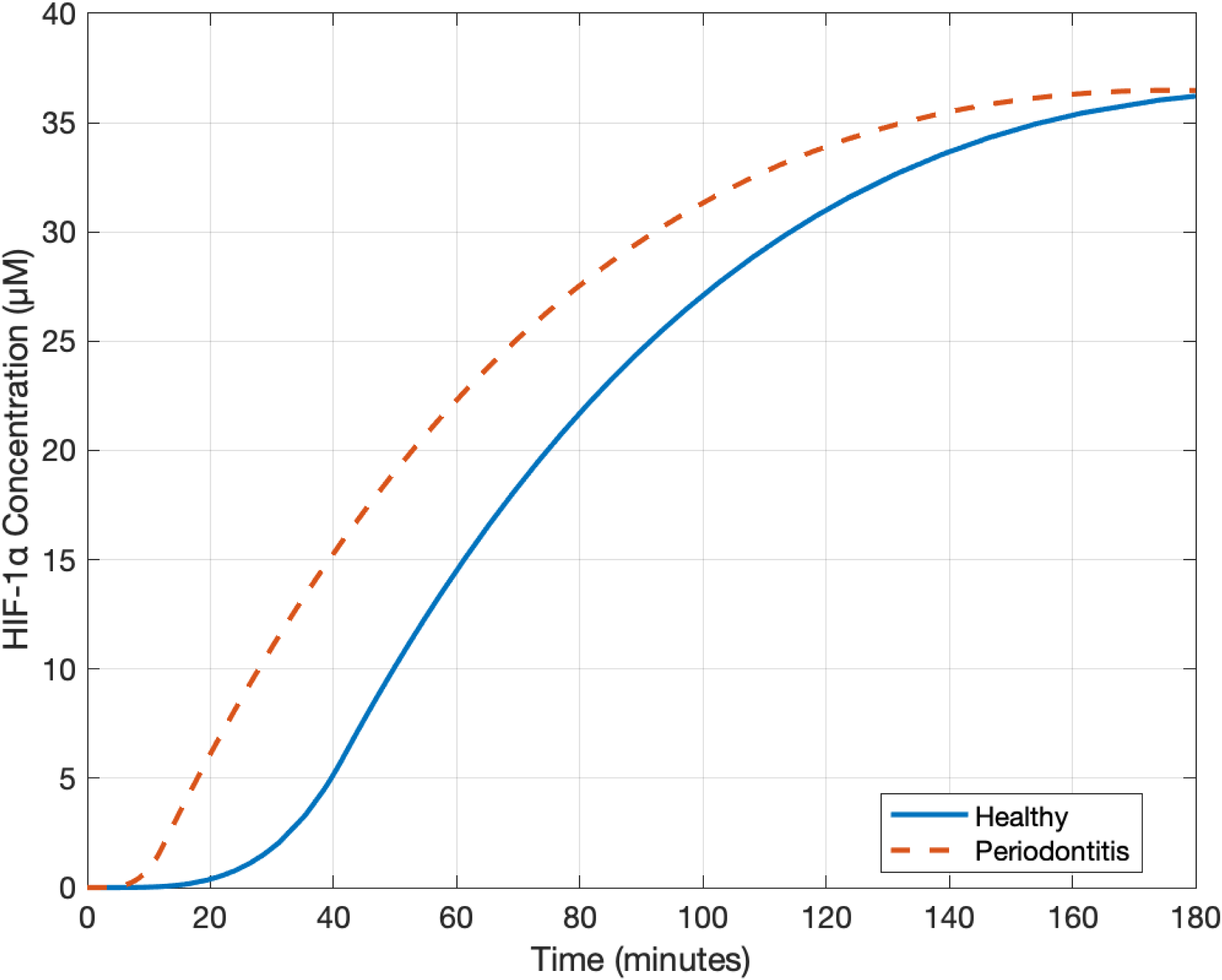
HIF-1α Concentration Dynamics Under Healthy and Periodontitis Conditions. Stabilization and activation of HIF-1α, strongly enhanced by lactate accumulation during chronic periodontal inflammation.

**Figure S10.**
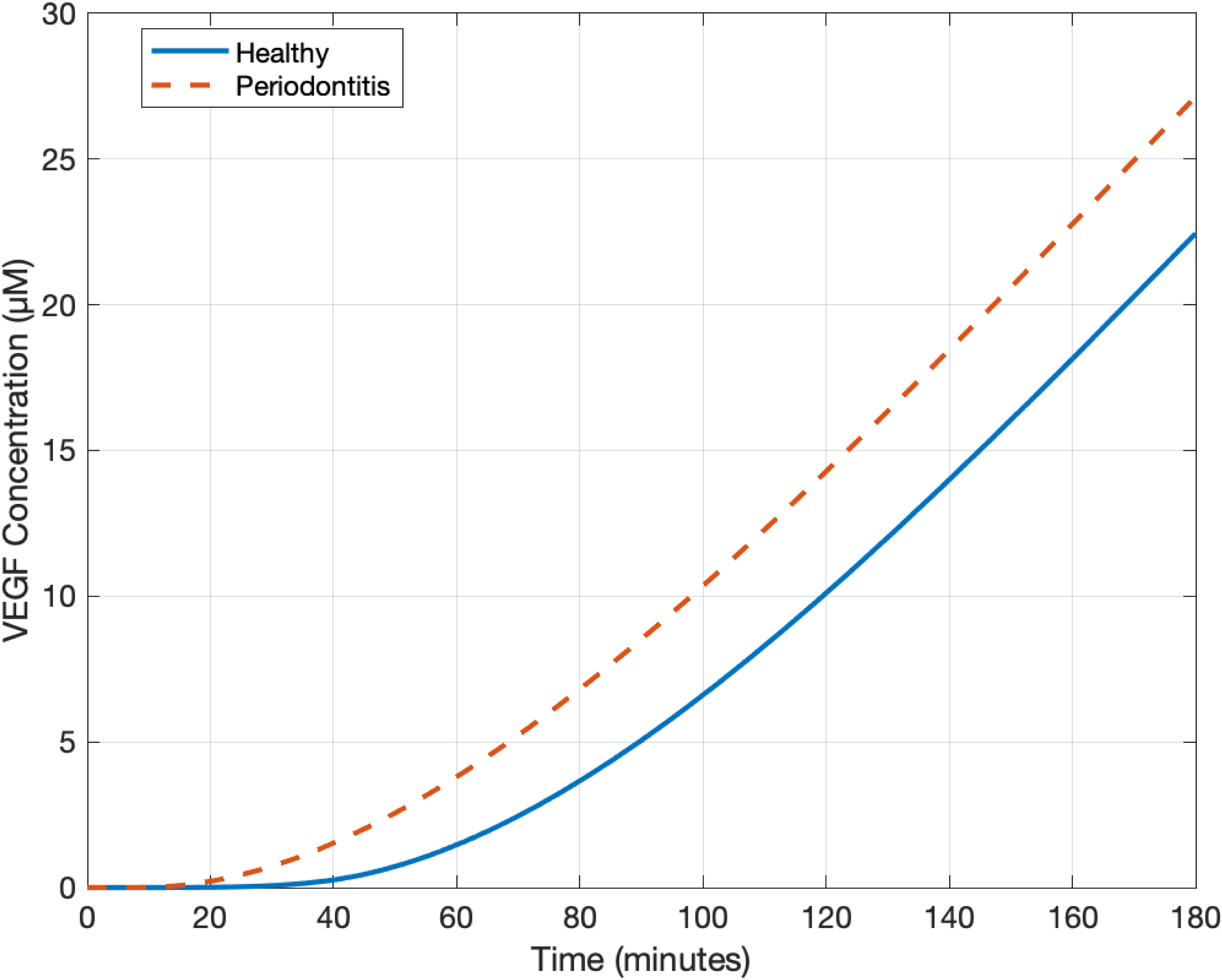
VEGF Concentration Dynamics Under Healthy and Periodontitis Conditions. VEGF expression kinetics demonstrating significantly increased angiogenic signaling induced by periodontal pathogen-mediated inflammation.

